# Injury-induced tau pathology promotes aggressive behavior in Drosophila without neurodegeneration

**DOI:** 10.1101/2025.11.22.689595

**Authors:** Roilea Maxson, Christine J. Smoyer, Megan F. Hampton, Yusheng Shen, Kailea Wiese, Cheryl Yee, Aishini Singh, Alexandria Funtila, Richard J. McKenney, Kassandra M. Ori-McKenney

## Abstract

The microtubule-associated protein tau is implicated in neurodegenerative diseases, but its physiological roles remain poorly understood. Here, we find that pan-neuronal expression of human tau (HsTau) in Drosophila coupled with injury triggers hyper-aggression in male flies, which is absent in flies expressing non-phosphorylatable tau. These behavioral manifestations result from activation of dopaminergic circuits without neurodegeneration. Using in vitro reconstitution assays, we find that phosphorylated HsTau maintains microtubule binding but loses its ability to suppress catastrophes, thereby promoting microtubule dynamicity. In contrast, unphosphorylated HsTau and fly tau (DmTau) stabilize microtubules by reducing catastrophe frequency. Our findings challenge the canonical view of tau as a simple microtubule stabilizer and instead position it as a dynamic regulator of microtubule function and neuronal excitability. These results reveal how acute tau phosphorylation can alter neural circuit function and behavior prior to neurodegeneration, providing new insights into tau’s physiological and pathological roles.

## INTRODUCTION

Traumatic brain injury (TBI) is a major public health concern and a leading risk factor for neurodegenerative diseases such as Alzheimer’s disease and chronic traumatic encephalopathy (CTE)^1–4^. Mild TBI, which accounts for 80-90% of cases^5^, frequently causes acute behavioral changes including increased aggression, impulsivity, and sleep disturbances^6,7^. These symptoms often precede cognitive decline and are associated with elevated levels of phosphorylated tau protein in patient serum, making tau a key molecular player in post-TBI pathology^8,9^.

Tau is a microtubule-associated protein (MAP) canonically described as a stabilizer of axonal microtubules, which are dynamic cytoskeletal polymers essential for neuronal structure, intracellular transport, and synaptic function^10–12^. MAPs can regulate microtubule dynamics by influencing their transition between growth and shrinkage phases, a process known as dynamic instability^13^. The prevailing hypothesis suggests that tau stabilizes microtubules by preventing this dynamicity, thereby maintaining overall polymer mass, but under neurodegenerative conditions, tau hyperphosphorylation reduces its microtubule binding affinity, leading to microtubule destabilization and impaired axonal transport^14,15^. This model has been challenged by a recent study proposing that tau may function to maintain microtubule dynamicity rather than simply enforcing stability, and that its loss might paradoxically lead to microtubule over-stabilization through other MAPs^16^.

Critical gaps remain in understanding not only the physiological function of tau, but how acute tau phosphorylation following injury affects neuronal function and behavior before neurodegeneration occurs^9,17^. To investigate these mechanisms, we employed a Drosophila model of injury, which offers powerful tools for dissecting molecular and circuit-level processes^18,19^. We demonstrate that injury induces phosphorylation of ectopically expressed HsTau within the fly brain resulting in hyper-aggressive behavior, which is dependent upon dopaminergic circuit activation. Using in vitro reconstitution assays with purified proteins, we find that phosphorylated HsTau maintains its microtubule binding but loses its ability to suppress catastrophes, thereby promoting microtubule dynamics. Our work reveals an acute, phosphorylation-dependent role for HsTau in modulating neuronal excitability and behavior following injury.

## RESULTS

### Pan-neuronal HsTau expression and injury lead to increased aggression

In this study, we utilized a previously established high-intensity trauma (HIT) device to induce injury in 6-8 day old virgin male flies expressing human 2N4R tau pan-neuronally via the Elav-GAL4 driver (Figure 1A-B)^18^. Although injury significantly increased mortality in Elav-GAL4>UAS-Tau flies compared to sham controls (6.93% vs. 0.95%; Figure S1a), overall lethality remained lower than previously reported rates^18^. Tau expression and injury have individually been shown to have pathophysiological effects on Drosophila^18,20,21^, as well as to cause behavioral deficits^22,23^. Therefore, we sought to observe fly behavior as a readout of tau pathology in response to injury. To assess general fly health, locomotion, and response to stimuli, we first used a negative geotaxis assay^24^. Surprisingly, while injury alone reduced climbing ability, flies expressing pan-neuronal HsTau showed a significant increase in climbing speed at 10 seconds post-startle in both sham and injury conditions compared to UAS-Tau controls (Figure 1C). This indicates that HsTau expression, even when combined with injury, does not lead to a general motor deficit, but rather to an unexpected increase in locomotor activity.

**Figure 1.**
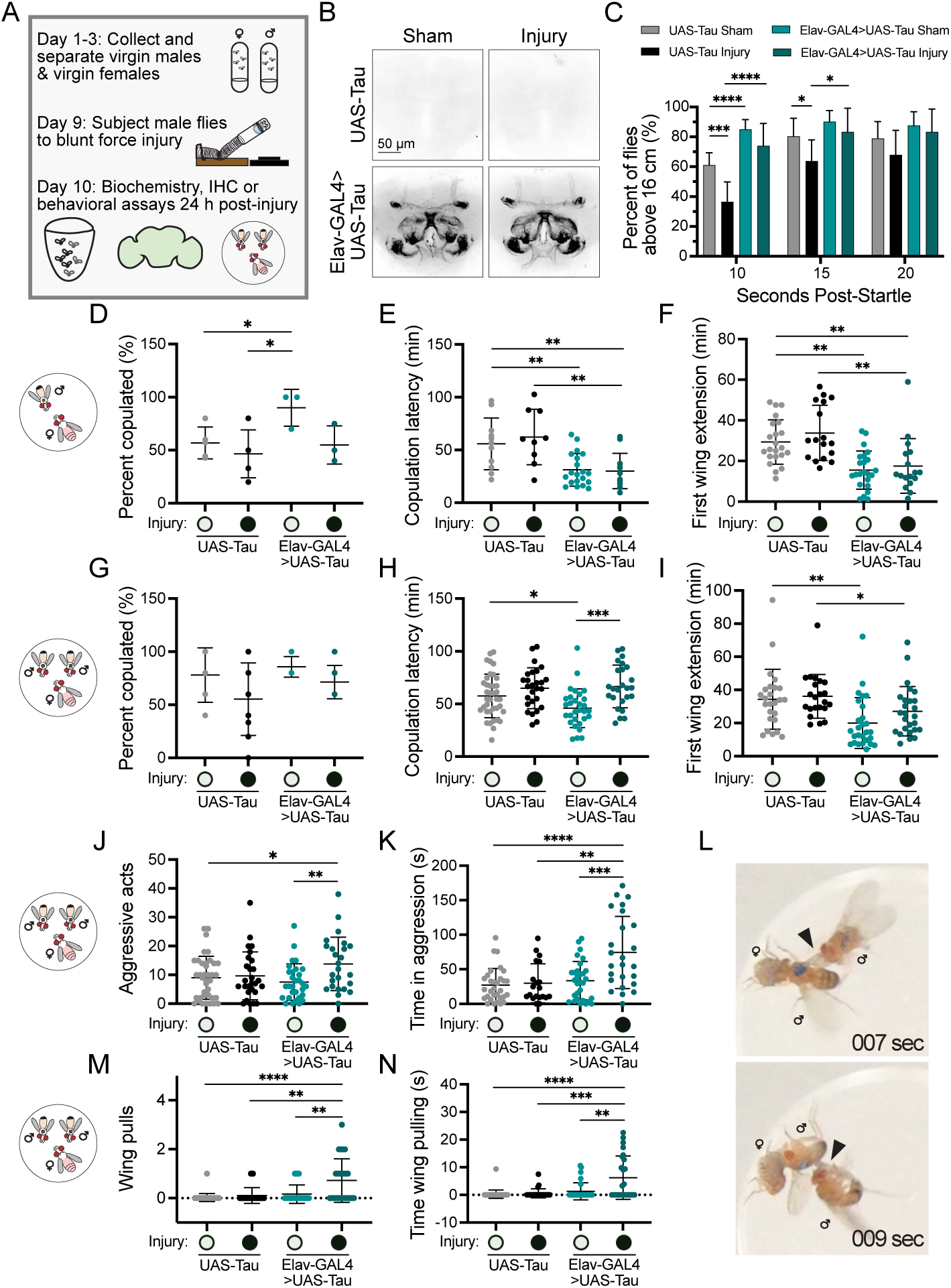
Pan-neuronal human tau Experimental setup for collection and administration of injury to *Drosophila* virgin males. Days indicate post-eclosion. All experiments were conducted at 24 hours post-injury. **B.** Representative max projection images of whole brain immunohistochemistry using an antibody against human tau in UAS-Tau (control) and Elav-GAL4>UAS-Tau male flies for sham and injury conditions. Scale bars: 50 μm. **C.** Quantification of the percent of male flies with the indicated genotype and injury condition that climb 16 cm within 10, 15, and 20 sec in a negative geotaxis assay (n=8 trials). For UAS-Tau sham *vs.* UAS-Tau injury at 10, 15, and 20 seconds post-startle: *p*=0.0010, 0.0252, and 0.2415, respectively; UAS-Tau sham vs. Elav-GAL4>UAS-Tau sham at 10, 15, and 20 seconds post-startle, *p*<0.0001, *p*=0.0756, and *p*=0.1179, respectively; UAS-Tau injury vs. Elav-GAL4>UAS-Tau injury at 10, 15, and 20 seconds post-startle, *p<*0.0001, 0.0219, and 0.0742, respectively; Elav-GAL4>UAS-Tau sham vs. Elav-GAL4>UAS-Tau injury at 10, 15, and 20 seconds post-startle, *p=*0.0769, 0.2951, and 0.5091, respectively. **D-F.** Noncompetitive courtship assays for UAS-Tau sham and injury male flies and Elav-GAL4>UAS-Tau sham and injury male flies (n=11, 9, 20, and 12 mating groups, respectively, from 3 independent trials). **D.** Quantification of the percent of fly pairs that mated within 2 hours. For UAS-Tau sham vs. UAS-Tau injury *p=*0.4212; UAS-Tau sham vs. Elav-GAL4>UAS-Tau sham *p=*0.0284; UAS-Tau injury vs. Elav-GAL4>UAS-Tau injury *p=*0.6060; Elav-GAL4>UAS-Tau sham vs. Elav-GAL4>UAS-Tau injury *p=*0.0724; UAS-Tau injury vs. Elav-GAL4>UAS-Tau sham *p=*0.0297. **E.** Quantification of latency time to copulation. For UAS-Tau sham vs. UAS-Tau injury *p=*0.5777; UAS-Tau sham vs. Elav-GAL4>UAS-Tau sham *p=*0.0018; UAS-Tau injury vs. Elav-GAL4>UAS-Tau injury *p=*0.0028; Elav-GAL4>UAS-Tau sham vs. Elav-GAL4>UAS-Tau injury *p=*0.8435; UAS-Tau sham vs. Elav-GAL4>UAS-Tau injury *p=*0.0074. **F.** Quantification of time to first wing extension by the male fly. UAS-Tau sham vs. UAS-Tau injury *p=*0.2640; UAS-Tau sham vs. Elav-GAL4>UAS-Tau sham *p*<0.0001; UAS-Tau injury vs. Elav-GAL4>UAS-Tau injury *p=*0.0014; Elav-GAL4>UAS-Tau sham vs. Elav-GAL4>UAS-Tau injury *p=*0.5784; UAS-Tau sham vs. Elav-GAL4>UAS-Tau injury *p=*0.0045. **G-N.** Competitive courtship assays for UAS-Tau sham and injury male flies and Elav-GAL4>UAS-Tau sham and injury male flies (n=38, 27, 30, and 25 groups, respectively, from 10 independent trials). **G.** Quantification of percent of flies that mated within 2 hours. For UAS-Tau sham vs. UAS-Tau injury *p=*0.1109; UAS-Tau sham vs. Elav-GAL4>UAS-Tau sham *p=*0.4648; UAS-Tau injury vs. Elav-GAL4>UAS-Tau injury *p=*0.2659; Elav-GAL4>UAS-Tau sham vs. Elav-GAL4>UAS-Tau injury *p=*0.0639. **H.** Quantification of latency time to copulation. For UAS-Tau sham vs. UAS-Tau injury *p=*0.1535; UAS-Tau sham vs. Elav-GAL4>UAS-Tau sham *p=*0.0175; UAS-Tau injury vs. Elav-GAL4>UAS-Tau injury *p=*0.7599; Elav-GAL4>UAS-Tau sham vs. Elav-GAL4>UAS-Tau injury *p=*0.0002. **I.** Quantification of time to first wing extension by the male fly. For UAS-Tau sham vs. UAS-Tau injury *p=*0.6975; UAS-Tau sham vs. Elav-GAL4>UAS-Tau sham *p=*0.0034; UAS-Tau injury vs. Elav-GAL4>UAS-Tau injury *p=*0.0321; Elav-GAL4>UAS-Tau sham vs. Elav-GAL4>UAS-Tau injury *p=*0.0942. **J.** Quantification of the total number of aggressive acts exhibited by male flies within the 10-minute window prior to copulation. For UAS-Tau sham vs. UAS-Tau injury *p=*0.7363; UAS-Tau sham vs. Elav-GAL4>UAS-Tau sham *p=*0.3929; UAS-Tau injury vs. Elav-GAL4>UAS-Tau injury *p=*0.0968; Elav-GAL4>UAS-Tau sham vs. Elav-GAL4>UAS-Tau injury *p=*0.0045; UAS-Tau sham vs. Elav-GAL4>UAS-Tau injury *p=*0.0271. **K.** Quantification of the time spent engaged in aggressive acts by male flies in the 10-minute window prior to copulation. UAS-Tau sham vs. UAS-Tau injury *p=*0.7200; UAS-Tau sham vs. Elav-GAL4>UAS-Tau sham *p=*0.3294; UAS-Tau injury vs. Elav-GAL4>UAS-Tau injury *p=*0.0013; Elav-GAL4>UAS-Tau sham vs. Elav-GAL4>UAS-Tau injury *p=*0.0005; UAS-Tau sham vs. Elav-GAL4>UAS-Tau injury *p<*0.0001. **L.** Representative images of a wing pull act. Genders are indicated. Arrowheads indicate wing being pulled. **M.** Quantification of the total number of wing pulls by male flies in the 10-minute window prior to copulation. For UAS-Tau sham vs. UAS-Tau injury *p=*0.1660; UAS-Tau sham vs. Elav-GAL4>UAS-Tau sham *p=*0.0434; UAS-Tau injury vs. Elav-GAL4>UAS-Tau injury *p=*0.0016; Elav-GAL4>UAS-Tau sham vs. Elav-GAL4>UAS-Tau injury *p=*0.0032; UAS-Tau sham vs. Elav-GAL4>UAS-Tau injury *p<*0.0001. **N.** Quantification of the time engaged in wing pulling by male flies in the 10-minute window prior to copulation. For UAS-Tau sham vs. UAS-Tau injury *p=*0.5070; UAS-Tau sham vs. Elav-GAL4>UAS-Tau sham *p=*0.0753; UAS-Tau injury vs. Elav-GAL4>UAS-Tau injury *p=*0.0005; Elav-GAL4>UAS-Tau sham vs. Elav-GAL4>UAS-Tau injury *p=*0.0025; UAS-Tau sham vs. Elav-GAL4>UAS-Tau injury *p<*0.0001. Two-sided unpaired Student’s *t*-tests were used to determine all *p*-values. For d-n graphs, all datapoints are plotted with lines indicating means ± s.d.

To investigate the effects of HsTau expression and injury on social behaviors, we performed non-competitive courtship assays between one virgin male and one virgin female and observed a significant reduction in copulation latency and the time to first male wing extension in tau-sham and tau-injury males compared to control-sham and control-injury males (Figure 1D-F). Next, we examined courtship and mating in a competitive courtship assay, which involves two males and one female (Figure 1G)^25^. Similar to the non-competitive assay, pan-neuronal HsTau expression led to a decrease in copulation latency and time to first wing extension in the absence of injury (Figure 1H-I). However, when coupled with injury, copulation was significantly delayed compared to tau-expressing males in the sham condition (66.6 vs. 46.0 min; Figure 1H).

To understand why injury coupled with HsTau expression caused a delay in copulation initiation in the competitive assay but not in the non-competitive assay, we examined inter-male aggression, which is known to occur when multiple virgin males are in the presence of a virgin female^26–28^. We observed a dramatic increase in aggressive behaviors for tau-injury compared to tau-sham males. Specifically, there was a nearly two-fold increase in both the total number of aggressive acts (12.6 vs. 6.9 acts) and the time spent engaged in aggression (74.3 vs. 33.6 seconds) for tau-injury compared to tau-sham males (Figure 1J-K). Intriguingly, tau-injury males also exhibited a unique aggressive behavior: wing-pulling (Figure 1L)^29^. Quantification revealed significantly more acts of wing pulling (Figure 1M) and a longer time spent in the act of wing pulling (Figure 1N) in tau-injury males compared to all other conditions. These results collectively indicate that pan-neuronal HsTau expression in combination with injury specifically promotes an increase in displays of aggressive behavior, including a unique aggressive act.

### Dopaminergic neuronal activation drives injury-induced tauopathy aggression

To identify the neuronal circuits responsible for the observed increase in aggression upon tau expression and injury, we performed a GAL4 driver screen using fly lines previously implicated in aggression (Figures 2 and S2)^30,31^. We drove human 2N4R tau expression in specific neuronal subtypes and assayed aggressive behavior in the competitive courtship setup (Figure 2A and S2A). We identified three GAL4 lines, Ddc-GAL4, 5HT7-GAL4, and Ple-GAL4, that, when driving HsTau expression coupled with injury, produced a significant increase in the total time spent in aggression (injury vs. sham treatment: Ddc 18.8 vs. 9.7 sec; 5HT7 14.9 vs.9.2 sec; Ple 17.5 vs. 7.2 sec), but only Ple-GAL4 driving tau expression coupled with injury significantly increased the number of aggressive acts compared to sham treatment (13.4 vs. 7.6 acts; Figure 2B-C). Dopa decarboxylase (Ddc) is a catalyst in the synthesis of both dopamine and serotonin^32^, 5HT7 is a serotonin receptor^33^, and pale (Ple) encodes a tyrosine hydroxylase which is the rate limiting step in the synthesis of dopamine^34^.

**Figure 2.**
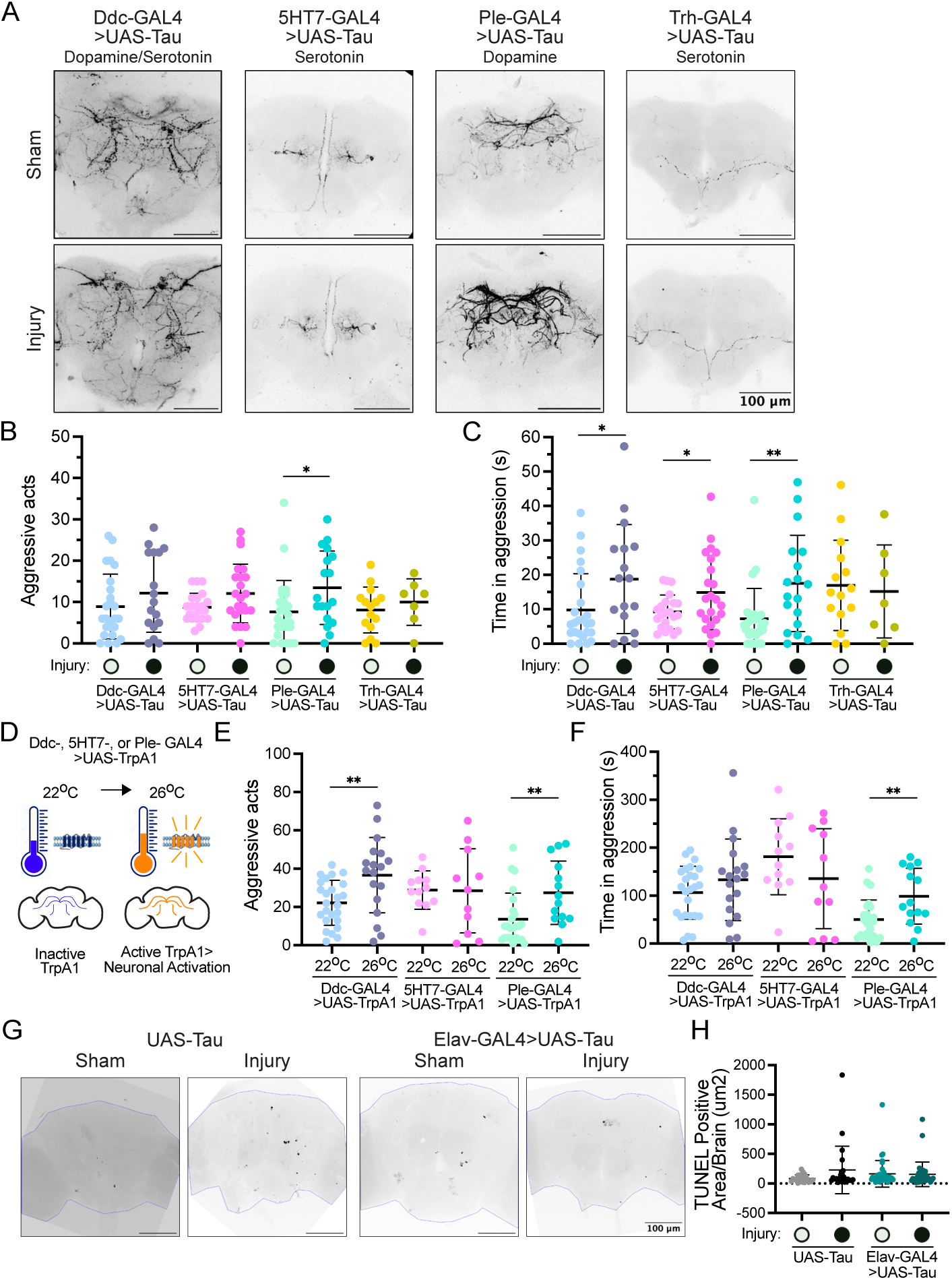
Human tau expression in dopaminergic neurons coupled with injury leads to neuronal activation and increased aggression in *Drosophila* male flies. **A.** Representative max projection images of whole brain immunohistochemistry using an antibody against human tau for Ddc-GAL4, 5HT7-GAL4, Ple-GAL4, or Trh-GAL4>UAS-Tau male flies for sham and injury conditions. Scale bars: 100 μm. **BC.** Competitive courtship assays for sham and injury male flies expressing UAS-Tau under Ddc-GAL4 (n=25 and 17 groups, respectively, from 8 independent trials), 5HT7-GAL4 (n=22 and 23 groups, respectively, from 7 independent trials), Ple-GAL4 (n=26 and 18 groups, respectively, from 9 independent trials), or Trh-GAL4 (n=15 and 7 groups, respectively, from 3 independent trials). Quantification for percent of flies that mated within 2 hours and latency time to copulation are in Extended Data Figure 3a-b. **B.** Quantification of total number of aggressive acts exhibited by male flies in the 10-minute window prior to copulation. Ddc-GAL4>UAS-Tau *p=*0.2322; 5HT7-GAL4>UAS-Tau *p=*0.0503; Ple-GAL4>UAS-Tau *p=*0.0243; Trh-GAL4 *p=*0.4566. **C.** Quantification of the time spent engaged in aggressive acts by male flies in the 10-minute window prior to copulation. Ddc-GAL4>UAS-Tau *p=*0.0328; 5HT7-GAL4>UAS-Tau *p=*0.0298; Ple-GAL4>UAS-Tau *p=*0.0045; Trh-GAL4 *p=*0.7751. **D.** Schematic of activating specific neuronal subtypes in male flies. Male flies expressing the temperature sensitive cation channel UAS-TrpA1 under Ddc-GAL4 (n=25 and 19 groups, respectively, from 4 independent trials), 5HT7-GAL4 (n=13 and 12 groups, respectively, from 2 independent trials), or Ple-GAL4 (n=26 and 15 groups, respectively, from 5 independent trials) were kept either at 26 °C to thermally activate specific neurons, or at 22 °C (no thermoactivation) for 1 hr. After 1 hr of thermal activation (or no activation), aggressive behavior was measured in a competitive courtship assay. Quantification for percent of flies that mated within 2 hours and latency time to copulation are in Extended Data Figure 3c-d. **E.** Quantification of total aggressive acts exhibited by male flies in the 10-minute window prior to copulation onset in a competitive courtship assay. Ddc-GAL4>UAS-TrpA1 *p=*0.0044; 5HT7-GAL4>UAS-TrpA1 *p=*0.9575; Ple-GAL4>UAS-TrpA1 *p=*0.0084. **F.** Quantification of total time spent in aggressive acts by male flies in the 10-minute window prior to copulation in a competitive courtship assay. Ddc-GAL4>UAS-TrpA1 *p=*0.2124; 5HT7-GAL4>UAS-TrpA1 *p=*0.2470; Ple-GAL4>UAS-TrpA1 *p=*0.0046. **G.** Representative max projection images of TUNEL staining in whole brains of UAS-Tau and Elav-GAL4>UAS-Tau male flies under sham or injury conditions. Blue outline indicates brain area. Scale bars, 100µm. **H.** Quantification of TUNEL positive area for each condition (n=22, 23, 38, and 36 brains). UAS-Tau sham vs. injury *p=*0.1039; Elav-GAL4>UAS-Tau sham vs. injury *p=*0.8935. Two-sided unpaired Student’s *t*-tests were used to determine *p*-values. All graphs display all datapoints with lines indicating means ± s.d.

To determine if these behaviors were due to neuronal activation or neuronal death, we employed the temperature-sensitive cation channel UAS-TrpA1 to activate specific neuronal subtypes (Figure 2D, schematic)^35^. We expressed UAS-TrpA1 under the control of Ddc-GAL4, 5HT7-GAL4, or Ple-GAL4 and maintained flies at either 22°C (inactive TrpA1) or 26°C (active TrpA1) for one hour. Activation of Ple-GAL4 controlled dopaminergic neurons was sufficient to produce a significant increase in both the total number of aggressive acts (27.4 vs. 13.6 acts; Figure 2E) and the time spent in aggression (98.7 vs. 50.0 sec; Figure 2F), mimicking the effects of HsTau expression coupled with injury. Attempts to silence Ple-GAL4 neurons using UAS-Kir resulted in a complete absence of copulation, precluding behavioral assessment (data not shown). Consistent with this, we found no evidence of neurodegeneration 24 hours post-injury upon pan-neuronal tau expression coupled with injury, as assessed by TUNEL staining for apoptotic cells (Figure 2G-H) and Western blot analysis of Dlg expression (Figure S3E-F). This suggests that the increased aggressive behavior is driven by neuronal activation, not neuronal death.

### HsTau phosphorylation is required for injury-induced aggression

We next investigated the molecular basis for this neuronal activation. Phosphorylation of tau is a critical factor in a range of pathologies including post-traumatic brain injury in humans^36,37^.

Using Elav-GAL4, we drove expression of UAS-Tau-S11A, where 11 commonly phosphorylated serines and threonines in human tau are mutated to alanines, rendering them non-phosphorylatable^38^. Pan-neuronal expression of tau-S11A did not lead to an increase in aggressive acts or time spent in aggression after injury (Figure 3A-B), indicating that tau phosphorylation is essential for the manifestation of these aggressive behaviors. Furthermore, we observed an increase in overall tau phosphorylation in fly head lysates 24 hours post-injury compared to sham controls (Figure 3C).

**Figure 3.**
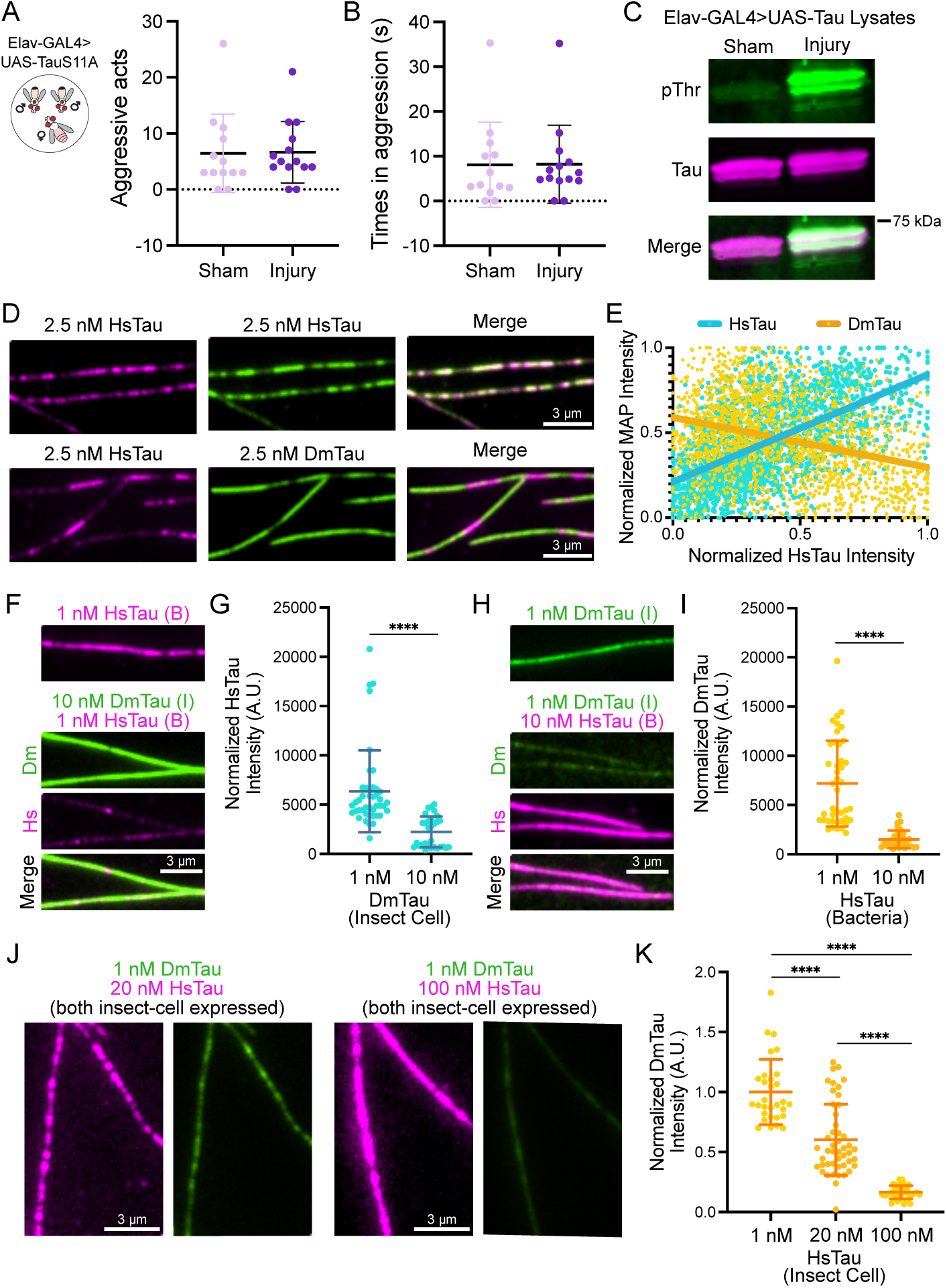
Human tau and fly tau compete for binding on microtubules. A-B. Competitive courtship assays for Elav-GAL4>UAS-TauS11A sham and injury flies (n=13 and 14 groups, respectively, from 3 independent trials). A. Quantification of the total number of aggressive acts exhibited by male flies within the 10-minute window prior to copulation. For Elav-GAL4>UAS-TauS11A sham vs. injury, *p=*0.9406. B. Quantification of the time spent engaged in aggressive acts by male flies in the 10-minute window prior to copulation. For Elav-GAL4>UAS-TauS11A sham vs. injury, *p=*0.9709. C. Western blot analysis of lysates generated from the heads of Elav-GAL4>UAS-Tau male flies subjected to sham or injury treatment. Blots were probed with antibodies against human tau (ms T46) and pan phospho-threonine (rb pThr). D. TIRF-M images of 2.5 nM mScarlet-2N4R-HsTau with 2.5 nM sfGFP-2N4R-HsTau show colocalization and miscibility on the microtubule, while 2.5 nM mScarlet-2N4R-HsTau and 2.5nM sfGFP-DmTau exclude each other into homotypic patches on the microtubule. Scale bars = 3 μm. E. Graph displaying individual xy pairs per pixel for mScarlet-HsTau intensity vs. sfGFP-HsTau or sfGFP-DmTau intensity on the microtubule, fit with a linear regression. Pearson’s correlation coefficients: 0.6080 for HsTau vs. HsTau (n=2042 XY pairs from n>15 microtubules, *p<*0.0001) and -0.3216 for HsTau vs. DmTau (n=1830 XY pairs from n > 15 microtubules, *p<*0.0001). F. TIRF-M images of 1 nM bacterially-expressed mScarlet-2N4R-HsTau in the absence or presence of 10 nM insect-cell expressed sfGFP-DmTau. Scale bars = 3 µm. **G.** Quantification of mScarlet-HsTau fluorescence intensity in the presence of 1 nM or 10 nM insect-cell expressed sfGFP-DmTau (n=42 and 30 microtubules, respectively from n=3 individual trials, *p<*0.0001. **H.** TIRF-M images of 1 nM insect-cell expressed sfGFP-DmTau in the absence or presence of 10 nM bacterially-expressed mScarlet-HsTau. Scale bars = 3 µm. **I.** Quantification of 1 nM sfGFP-DmTau normalized fluorescence intensity in the presence of 1 nM or 10 nM bacterially-expressed mScarlet-HsTau (n=42 and 30 microtubules, respectively from n=3 individual trials, *p<*0.0001). **J.** TIRF-M images of 1 nM insect-cell expressed sfGFP-DmTau in the presence of 20 nM or 100 nM insect-cell expressed mScarlet-HsTau. Scale bars = 3 µm. **K.** Quantification of 1 nM sfGFP-DmTau normalized fluorescence intensity in the presence of 1 nM, 20 nM or 100 nM insect-cell expressed mScarlet-HsTau (n=30, 47 and 33 microtubules, respectively from n=2 individual trials, **** indicates *p<*0.0001). (I = insect cell expression and B = bacterial expression). Two-sided unpaired Student’s *t*-tests were used to determine *p*-values. All graphs display all datapoints with lines indicating means ± s.d.

To understand the process by which tau phosphorylation post-injury leads to neuronal activation, we first investigated the binding behaviors of HsTau and DmTau on microtubules. We purified non-phosphorylated human 2N4R tau (HsTau expressed in bacteria) and phosphorylated human 2N4R tau (HsTau expressed in insect cells, mimicking its post-injury state), for comparison with fly tau (DmTau expressed in insect cells to reflect its native environment)(Figure S4A-C). Using total internal reflection fluorescence microscopy (TIRF-M), we imaged bacterially-expressed mScarlet-HsTau with either bacterially-expressed sfGFP-HsTau or insect cell-expressed sfGFP-DmTau on taxol-stabilized microtubules. HsTau is highly miscible and co-assembles into oligomeric assemblies on the microtubule lattice, which can be visualized when adding equimolar amounts of sfGFP-HsTau and mScarlet-HsTau (Figure 3D-E)^39^. In contrast, at equimolar concentrations, mScarlet-HsTau and sfGFP-DmTau exhibit negatively correlated binding behaviors, segregating into homotypic patches on the microtubule, which indicates competitive binding (Figure 3D-E)^40^. In support of this competition, we found that a 10-fold excess of DmTau displaces HsTau from the microtubule (Figure 3F-G), and similarly, a 10-fold excess of HsTau prevents DmTau from binding the lattice (Figure 3H-I). Even when HsTau is phosphorylated, HsTau and DmTau compete with one another for microtubule binding. When HsTau is expressed in insect cells and phosphorylated, its microtubule binding affinity is reduced^41,42^, but it still prevents the binding of DmTau when in excess (Figure 3J-K). These data suggest that overexpressed HsTau likely outcompetes endogenous DmTau from the microtubule lattice within fly neurons, and even when phosphorylated post-injury, HsTau likely maintains its dominant binding due to its excess concentration. In addition, these results reveal that DmTau is a molecularly distinct MAP from HsTau with respect to microtubule binding, consistent with a prior analysis of tau evolution^43^. These results provide a possible explanation for why ectopic expression of HsTau is necessary for injury-induced neuronal activation.

### Phosphorylation regulates the ability of HsTau to stabilize microtubules

To molecularly define how phosphorylated HsTau differs from DmTau to cause neuronal activation, we examined the effects of bacterially expressed HsTau, insect-cell expressed HsTau, and DmTau on microtubule dynamics, which are essential for neuronal activity and synaptic transmission (Figure 4A) ^44–46^. We employed an in vitro assay to reconstitute microtubule dynamics in which we affix stable GMPCPP microtubule seeds to the coverslip, then image as free tubulin in solution polymerizes from these seeds in the absence or presence of tau proteins (Figure 4A-B and S4D)^47^. We found that neither bacterial nor insect-cell HsTau affected microtubule growth rate, unlike DmTau which increased it (tubulin alone vs. DmTau: 0.72 vs. 0.99 µm/min; Figure 4C). We next asked how each tau protein regulates microtubule stability, which we define here as the suppression of catastrophe events, which is the transition from growth to shrinkage. Interestingly, phosphorylated insect-cell HsTau did not significantly alter catastrophe frequency, but both DmTau and bacterial non-phosphorylated HsTau significantly reduced catastrophes (tubulin alone vs. bacterial HsTau vs. DmTau: 0.23 vs. 0.18 vs. 0.06 cat/min; Figure 4D). All three forms of tau significantly increased rescue frequency though to different degrees (tubulin alone vs. bacterial HsTau vs. insect cell HsTau vs. DmTau: 0.036 vs. 0.200 vs. 0.098 vs. 0.156; Figure 4E). We observed similar effects in BEAS-2B cells expressing EB1-tdTomato, a microtubule plus-end binding protein that tracks growing microtubule ends and serves as a readout for polymerization dynamics. While HsTau expression maintained normal microtubule dynamics, DmTau expression promoted microtubule stability as evidenced by a decrease in microtubule polymerization events compared to GFP expression alone (GFP vs. DmTau expression: 591.7 vs. 375.3 events/min; Figure 4F-G). Neither HsTau or DmTau affected the dwell time of EB1 at the plus end, confirming these results were not due to an effect on the EB1 protein (Figure 4H-I). These results indicate that while DmTau and non-phosphorylated HsTau stabilize microtubules and promote growth by inhibiting catastrophes, phosphorylated HsTau does not, which enables increased microtubule dynamics. Therefore, we hypothesize that upon overexpression, HsTau initially outcompetes DmTau and potentially other fly MAPs, but as it becomes phosphorylated post-injury, its altered properties permit increased microtubule dynamicity, stimulating neuronal activity and contributing to the manifestation of aggressive behaviors observed in male flies.

**Figure 4.**
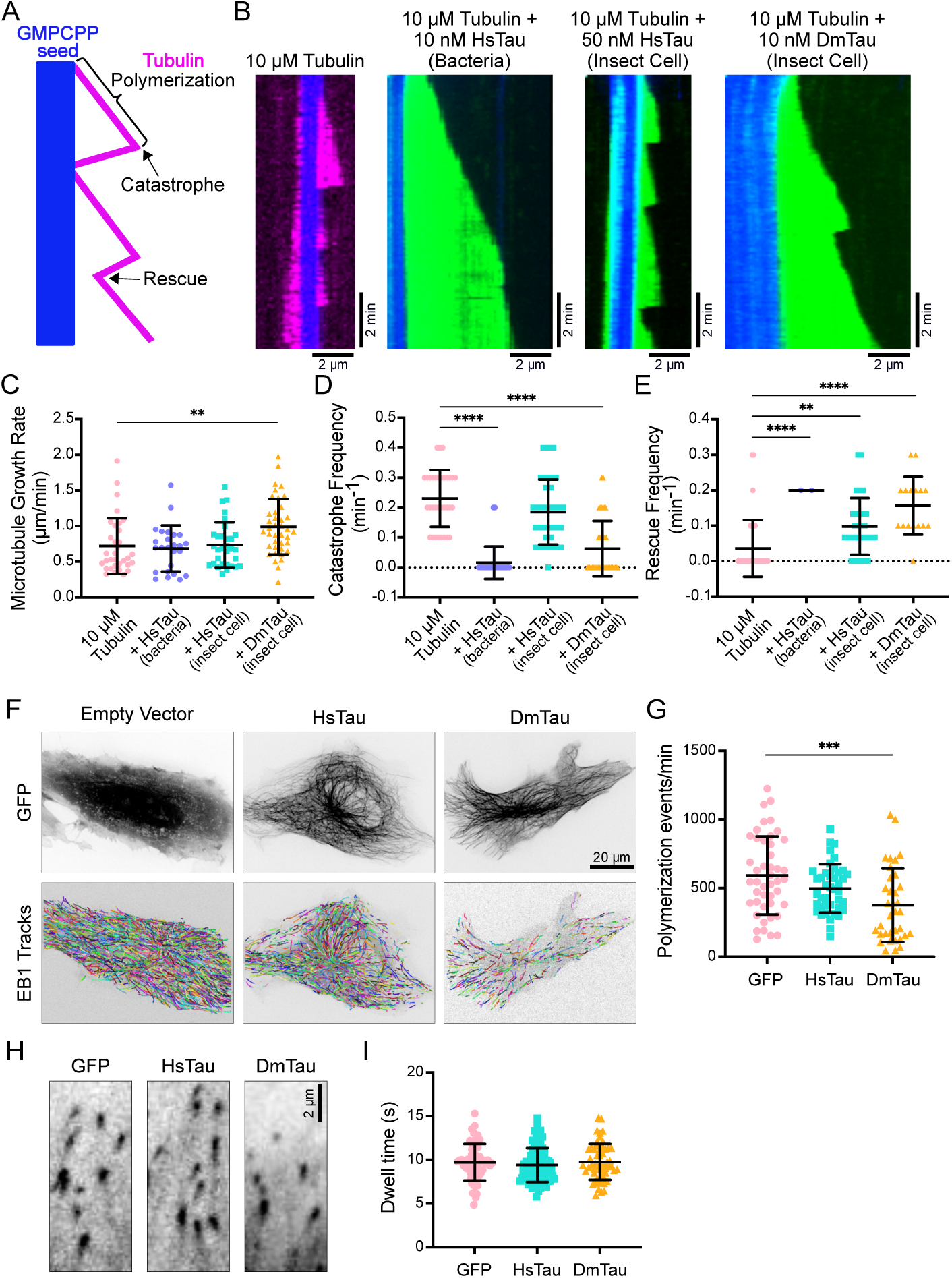
Human and fly tau exhibit distinct effects on microtubule dynamics in vitro and in cells. **A.** Schematic depicting an in vitro reconstituted microtubule dynamics assay with labeled parameters that we measured. A diagonal line represents tubulin (magenta) polymerizing off of a stable GMPCPP seed (blue), then undergoing a catastrophe and rapidly depolymerizing back to the GMPCPP seed. If the microtubule begins to polymerize again before reaching the GMPCPP seed, this is considered a rescue event. **B.** Representative kymographs of microtubule dynamics showing the polymerization of 10 μM tubulin (magenta) + 1 mM GTP from GMPCPP seeds (blue) in the absence or presence of human sfGFP-2N4R-tau (HsTau, green) expressed in bacteria or insect cells or *Drosophila melanogaster* sfGFP-tau (DmTau, green) expressed in insect cells at the indicated concentrations. Scale bars: y, 2 min; x, 2 μm. **C-E** Quantification of microtubule plus end growth rate (**C**), catastrophe frequency (**D**), and rescue frequency (**E**) for 10 μM tubulin + 1 mM GTP in the absence or presence of 10 nM bacterially-expressed HsTau, 50 nM insect cellexpressed HsTau, or 10 nM insect cell expressed-DmTau (n=33, 25, 31, and 39 analyzed kymographs from n=3 independent trials). For microtubule growth rate (**C**), tubulin alone vs. bacterial HsTau, *p=*0.7113; vs. insect-cell HsTau, *p=*0.8525; vs. DmTau, *p=*0.0048. For catastrophe frequency (**D**), tubulin alone vs. bacterial HsTau, *p<*0.0001; vs. insect-cell HsTau, *p=*0.0803; vs. DmTau, *p<*0.0001. For rescue frequency (**E**), tubulin alone vs. bacterial HsTau, *p<*0.0001; vs. insect-cell HsTau, *p=*0.0027; vs. DmTau, *p<*0.0001. **F.** Images of BEAS-2B cells expressing EB1-tdTomato in conjunction with either GFP empty vector, GFP-HsTau, or GFP-DmTau visualized by spinning disk confocal microscopy, with associated EB1-tdTomato comet trajectories represented by colored lines (2.5 fps for 3 min) showing the growth pattern of microtubules under each transfection condition. Scale bar: 20 µm. **G.** Quantification of polymerization events per minute for each transfection condition, GFP empty vector, GFP-HsTau, and GFP-DmTau (n=45, 41, and 35 cells, respectively from 3 independent experiments). For GFP vs. HsTau, *p=*0.0667; vs. DmTau, *p=*0.0008. **H.** Magnified view of EB1 comets under each transfection condition, GFP empty vector, GFP-HsTau, or GFP-DmTau. Scale bar: 2 µm. **I.** Quantification of EB1 dwell time under each transfection condition, GFP empty vector, GFP-HsTau, or GFP-DmTau (n=63, 85, and 56 cells, respectively). For GFP vs. HsTau, *p=*0.3398; vs. DmTau, *p=*0.9308. Two-sided unpaired Student’s *t*-tests were used to determine *p*-values. All graphs display all datapoints with lines indicating means ± s.d.

## DISCUSSION

Here, we use a *Drosophila* model of injury coupled with human tau expression to investigate the acute behavioral and neuronal consequences of injury. Our findings reveal that injury in the presence of HsTau, but not the non-phosphorylatable mutant Tau-S11A, rapidly induces hyperactive and aggressive behaviors within 24 hours. This behavioral shift is not a product of general motor deficits or neurodegeneration but is driven by the activation of dopaminergic neuronal circuits, as mimicked by direct thermogenetic stimulation. These results highlight a critical, phosphorylation-dependent role for tau in modulating neuronal activity and behavior immediately following injury, before the onset of any detectable neurodegenerative pathology.

This work challenges two long-standing paradigms in the tau field. First, it demonstrates that tau-dependent pathological changes can manifest as altered neuronal activity and behavior on a rapid timescale, preceding the classical hallmark of aggregation and neurodegeneration. This is highly relevant to the human condition, where traumatic brain injury patients frequently experience acute behavioral changes such as impulsivity, aggression, and sleep disturbances, which can persist and predict negative long-term outcomes^5,6,48^. Our model provides a system to dissect the molecular and circuit-based mechanisms underlying these acute post-injury symptoms.

Second, our data fundamentally reshape the understanding of tau’s core biochemical function. Contrary to the canonical view of tau as a microtubule stabilizer, our in vitro assays demonstrate the functional outcome of tau binding is context-dependent. DmTau and unphosphorylated HsTau suppress catastrophes and promote microtubule stability. Despite maintaining its ability to bind the microtubule lattice, phosphorylated HsTau (mimicking its post-injury state) loses its capacity to suppress catastrophes, thereby permitting enhanced microtubule dynamics. This suggests that the primary role of tau, particularly in its physiological state, may not be to statically stabilize microtubules but to maintain a requisite level of dynamicity. Our results provide a molecular explanation for prior work demonstrating that tau is enriched on the labile domains of axonal microtubules and its depletion leads to microtubule over-stabilization via the expansion of genuine microtubule stabilizers^16^, and are also consistent with in vivo observations of tau’s rapid exchange on microtubules and in vitro observations of tau’s avoidance of the GTP cap^49,50^. This revised model challenges the traditional "loss-of-function" hypothesis in neurodegeneration. Instead of causing destabilization, tau loss may allow other MAPs to dominate the lattice, resulting in excessive microtubule stability and reduced dynamics. An overly stable cytoskeleton could impair organelle transport and synpatic plasticity, inducing a state of premature cellular aging and functional decline before overt cell death. Thus, tau’s role may be to temper over-stabilization, ensuring the cytoskeletal flexibility necessary for healthy neuronal function.

Overall, our study positions tau not merely as a passive stabilizer lost in disease, but as a dynamic regulator of microtubule function and neuronal excitability. The acute behavioral phenotypes we observe following injury are likely a consequence of tau phosphorylation and subsequent changes in microtubule dynamics within specific neurochemical circuits. This model provides a unique opportunity to explore early molecular events following injury and could provide new therapeutic avenues to mitigate acute symptoms and potentially prevent long-term neurodegeneration.

## METHODS

### Fly stocks and husbandry

*Drosophila melanogaster* were raised on standard Bloomington cornmeal medium (Genesee Scientific 66-121) and maintained at 24°C under a 12-h light:12-h dark cycle. The following stock lines were used: UAS-Tau.wt.7B (BDSC_51363), elav-GAL4 (BDSC_49226); Ddc-GAL4 (BDSC_7009), 5HT7-GAL4 (BDSC_49414), Ple-GAL4 (BDSC_8848), Trh-GAL4 (BDSC_38388), UAS-TrpA1 (BDSC_26263), Fru-GAL4 (BDSC_30027), NPF-GAL4 (BDSC_25681), 5HT1A-GAL4 (BDSC_50443), Vsx2-GAL4 (BDSC_39599), Ct-GAL4 (BDSC_49808), UAS-Kir2.1 (BDSC_6595), and UAS-TauS11A (BDSC_51366). The UAS-TauS11A construct is the human 2N4R tau that has the following sites mutated to alanines so they cannot be phosphorylated: S46, T50, S199, T202, T205, T212, S214, T231, S235, S396 and S404). Flies used were F1 progeny of indicated GAL4 and UAS lines.

### Injury Administration

Virgin male and female flies were collected and housed separately until they were 6-8 days post-eclosion. Adult male flies were injured using the high-intensity trauma (HIT) device as previously described^18^. 20 flies were loaded into the HIT device, and received 4 hits, 5 minutes apart. For the sham condition, a vial of control flies was also loaded into vials for the same period of time, but received no hits. Flies were allowed to recover for 40 minutes before returning to food vials for 24 hours before subsequent assays.

### Brain Immunohistochemistry

Fly brains were dissected into 1X PBS+0.1% triton-X, fixed in 4% PFA+0.1% triton-X for 20 minutes at room temperature, and washed 3 times for 20 minutes. Samples were blocked in 5% normal goat serum (1X PBS+0.1% triton-X) for 30 minutes at room temperature, incubated with anti-Tau (1:500 T46, Invitrogen) primary antibody for 48 hours at 4°C, washed 3 times with 1X PBS+0.1% triton-X, incubated for 2 hours with Alexa Fluor-488 (1:500 anti-mouse) secondary antibody, and washed 3 times with 1X PBS+0.1% triton-X. Brains were mounted in Vectashield Antifade Mounting Media. Brains were imaged using a Zeiss LSM 980-Airyscan2.

For assessing neurodegeneration, the DeadEnd TUNEL System (Promega G3250) was used. Whole flies were fixed for 3 hours in 4% PFA (1X PBS + 0.1% triton-X) at room temperature, rinsed with 1X PBS+0.5% triton-X, dissected in cold 1X PBS+0.1% triton-X, and washed 3 times for 20 minutes in 1X PBS+0.1% triton-X. Brains were equilibrated in equilibration buffer for 10-minutes at room temperature, incubated with rTDT enzyme buffer (45ul equilibration buffer, 5ul nucleotide mix, 1ul rTDT enzyme, 1:1000 DAPI) overnight at 4°C. The reaction was stopped with 2X SSC for 1 minute, then incubated with fresh 2X SSC for 10-minutes. Brains were washed 3 times with 1X PBS+0.1% triton-X for 10-minutes at room temperature. All incubations and washes were performed in the dark. Samples were mounted in Vectashield Antifade Mounting Media and imaged using a Zeiss LSM 980-Airyscan2. Images were analyzed using ImageJ (Fiji). Max projection images were processed using the Autolocal Threshold with the Phansalkar method. Particles were counted with a lower size threshold of 1.5um, circularity 0-1. Results are displayed as total TUNEL positive area per brain.

### Negative Geotaxis Assay

Groups of 20 male flies were placed in 100 ml graduated cylinders marked at 16cm from the bottom. Flies were allowed to acclimate to cylinders for 10-minutes. For each trial, flies were tapped down and allowed to rest for 3 minutes. Each group of flies was recorded for 4 replicate trials. The number of flies above the 16cm mark was recorded for 10-, 15-, and 20-seconds post-startle.

### Courtship Assays

For non-competitive courtship assays, single virgin male and virgin female flies were loaded into plexiglass chambers (10mm diameter x 5 mm depth). Pairs were filmed for 2 hours at 24°C at consistent time of day. Copulation percentage represents pairs which mated within the 2 hours of filming. Copulation latency represents the time to successful copulation (determined by sustained contact during which the male mounts the female), wing-extension is characterized by approximately 90-degree extension of the wing while the male faces the female^51^. For competitive courtship assays, groups consisted of two virgin males and a single virgin female.

Aggressive acts between males were scored within the 10-minutes prior to the start of copulation for each group, and included fencing, boxing, lunging, tussling, head butting, shoving, holding, abdomen bending, wing pulling, and chasing. Total numbers of acts were scored and the total amount of time that male flies engaged in these aggressive acts were scored. For TrpA1 temperature sensitive activation, flies were raised at 22°C. For courtship assays, neurons were activated by acclimating flies to 26°C 1 hour before filming; genotype matched control flies were filmed at 22°C.

### Molecular Biology

cDNA for full-length human tau was previously purchased from Addgene (16316). cDNA for *Drosophila* tau was purchased from the Drosophila Genomics Resource Center (RE16764). For bacterial cell expression full length human tau (HsTau) was cloned into a pET28A vector via Gibson assembly. For insect-cell expression, HsTau and DmTau were cloned into pFastBac vectors via Gibson assembly. All pFastBac and pET28A tau constructs contain an N-terminal cassette consisting of a 6× His-tag, tandem Strep-tags, and fluorophore connected by a GS-linker as previously described (Tan et al., 2019). The pFastBac and pET28 sfGFP-2N4R-HsTau produce the exact same amino acid sequence. For Beas-2B cell expression, HsTau and DmTau were cloned into a pEGFP-C1 vector and we used EB1-tdTomato (Addgene plasmid 50825). All constructs were verified by DNA sequencing (Plasmidsaurus).

### Protein expression and purification

Bacterially expressed HsTau was purified from BL21 cells as previously described^39^. BL21 cells were grown at 37°C until an optical density at 600nm of 0.5 was reached. Protein expression was then induced with 0.4mM isopropyl-β-D-thiogalactoside overnight at 24°C, cells were harvested and frozen at -80°C. Cell pellets were resuspended in lysis buffer (50 mM MES (pH 6.0), 10 mM EDTA, 10 mM DTT supplemented with 1mM DTT, 0.2mM PMSF, and 1:1000 protease inhibitor mix (Promega, Madison, WI). Cell lysis was performed using an Emulsiflex C-3 (Avestin), 5μl of nuclease were added into per 40 ml lysed cells. Lysed cells were centrifuge at 20,000xg for 20 minutes at 4°C. Clarified lysate was diluted 5x, filtered, and purified via cation exchange chromatography using a HiTrap S column (Cytiva). The protein was then eluted with a linear gradient of NaCl from 0 to 1 M. Protein-containing fractions were pooled and precipitated using 0.3 g/ml ammonium sulfate and incubated 30 min at 4°C. Precipitated proteins were then centrifuged at 20,000 x g for 30 min at 4°C, and resuspended in 10 mM phosphate buffer (PB), pH 7.4, with 10 mM DTT. Resuspended proteins were further purified by Superdex 200 10/300 size exclusion column in 10 mM PB. Fractions were collected and analyzed by SDS-PAGE and stained with Coomassie blue. Protein-containing fractions were pooled and concentrated to higher than 2 mg/ml using molecular weight concentrators with a cut-off filter of 3 kDa (MilliporeSigma). For baculovirus expression of HsTau and DmTau, the Bac-to-Bac protocol (Invitrogen) was followed. SF9 cells were grown in shaker flasks to ∼2x10⁶/ml and infected at a ratio of 1:100ml virus to cells. The infection was allowed to proceed for 48-65 hours before cells were harvested and frozen in liquid nitrogen. Cell pellets were resuspended in lysis buffer (50mM Tris-HCl pH 8.0 150 mM K-acetate, 2 mM Mg-acetate, 1 mM EGTA supplemented with 1mM DTT, 0.2mM PMSF, and 1:1000 protease inhibitor mix (Promega, Madison, WI). Cells were lysed with a dounce homogenizer and centrifuged at 14,000xg for 20 minutes at 4°C. Clarified lysate from either bacterial or baculovirus expression was passed over a column with Streptactin Superflow resin (Qiagen). After incubation, the column was washed with lysis buffer, and purified proteins eluted with lysis buffer supplemented with 50mM Biotin. Proteins were loaded onto a HiTrap SP HP column and purified via ion exchange and eluted into a buffer containing 50mM Tris pH 7.5, 1mM EGTA, 2mM MgSO4 over a gradient of 0 to 0.6M NaCl.

Protein containing fractions were pooled and concentrated using a 50kDa Amicon centrifugal concentrator (MilliporeSigma), and frozen at -80°C. Insect-cell HsTau has been previously validated to be phosphorylated by mass spec and PhosTag gel^41^, but bacterially expressed and insect-cell expressed HsTau were validated as non-phosphorylated and phosphorylated by running samples on an SDS-PAGE gel and staining with Pro-Q™ Diamond Phosphoprotein Gel Stain (ThermoFisher).

### Preparation of Microtubules

For microtubules, porcine brain tubulin was isolated using a high-molarity Pipes procedure and then labeled with biotin NHS ester, Dylight-405 NHS ester, or Alexa647 NHS ester, as described previously (https://mitchison.hms.harvard.edu/files/mitchisonlab/files/labeling_tubulin_and_quantifying_la beling_stoichiometry.pdf). A mixture of brain tubulin, biotin-tubulin, and 405-fluorescent-tubulin purified from porcine brain (∼10:1:1 ratio) was assembled in BRB80 buffer (80 mM Pipes, 1 mM MgCl2, 1 mM EGTA, pH 6.8 with KOH) with 1 mM GTP for 20 minutes at 37°C, and then polymerized microtubules were stabilized with 20 µM taxol and incubated for an additional 30 min at 37°C. Microtubules were pelleted over a 25% sucrose cushion in BRB80 buffer supplemented with 5 µM taxol to remove unpolymerized tubulin, and the pellets were resuspended in BRB80 with 10 µM taxol. GMPCPP microtubules were polymerized by combining unlabeled tubulin, 650-tubulin, and biotin-tubulin (∼10:1:1) to a concentration of ∼60mM in BRB80 supplemented with 1mM DTT and 1mM GMPCPP seed (Jena BioScience NU-405S), then incubating at 37°C for 20-30 minutes. Microtubules were pelleted over a 25% sucrose cushion and resuspended in BRB80 supplemented with 1mM DTT.

### TIRF microscopy

TIRF microscopy experiments were performed on an inverted research microscope Eclipse Ti2-E with the Perfect Focus System (Nikon), equipped with a 1.49 NA 100× TIRF objective with the 1.5× tube lens setting, a Ti-S-E motorized stage, piezo Z-control (Physik Instrumente), LU-N4 laser units (Nikon) as the light source, an iXon DU897 cooled EMCCD camera (Andor) with an high-speed emission filter wheel (ET480/40M for mTurquoise2, ET525/50M for GFP, ET520/40M for YFP, and ET632/60M for mRuby2; Chroma). The microscope was controlled with NIS Elements software (Nikon). TIRF chambers were assembled by combining an acid-washed glass coverslip prepared as previously described (http://labs.bio.unc.edu/Salmon/protocolscoverslippreps.html), a pre-cleaned slide (Fisher Scientific), and double-sided sticky tape. The chambers were first incubated with 0.5 mg/ml PLL-PEG-biotin (Surface Solutions Group. Chicago, IL) for 5-10-minutes, followed by 0.5 mg/ml streptavidin. Microtubules were diluted in BRB80 supplemented with 10 μM taxol and added into streptavidin-adsorbed flow chambers and incubated for 2 minutes at room temperature. Chambers were subsequently washed with SRP90 assay buffer (90 mM HEPES, 50 mM KCH3COO, 2 mM Mg(CH3COO)2, 1 mM EGTA, 10% glycerol, pH = 7.6), supplemented with 1 mg/mL BSA, 0.05 mg/mL biotin–BSA, 0.1 mg/mL K-casein, 0.5% Pluronic F-127, and 10 μM taxol). Purified proteins were diluted to indicated concentrations in the assay buffer and flowed into the TIRF chamber. Data were analyzed manually using ImageJ (Fiji).

For MT dynamics assay GMPCPP microtubules were diluted in 1X BRB80, added into streptavidin-absorbed flow chambers, and incubated for 2 minutes at room temperature, followed by BRB80 assay buffer (BRB80, 1 mg/mL BSA, 0.05 mg/mL biotin–BSA, 0.2 mg/mL K-casein, 0.5% Pluronic F-127) supplemented with 2mM GTP, unlabeled tubulin, 405-labeled tubulin (concentration and ratios indicated), PCA, PCD, Trolox, and indicated concentration of protein (diluted in BRB80 assay buffer). Protein concentrations were determined based on prior work^39,41^. The reaction chamber was mounted on a pre-warmed microscope objective to 30°C by a temperature collar. Microtubule seeds and the dynamic ends were excited at 650 and 405 nm, respectively. The movies were captured within 5 seconds/frame for 10-20 minutes.

### Live cell imaging

BEAS-2B cells (ATCC, CRL-9609) were maintained in Dulbecco’s modified Eagle’s medium (DMEM, Gibco), supplemented with 10% fetal bovine serum (FBS), 50 units/mL of penicillin and 50 μg/mL of streptomycin. Cell cultures were maintained in a 95% air/5% CO2 atmosphere at 37°C. For live-cell imaging, cells were seeded at a density of approximately 5×10^4^ cm ^-2^ on a glass coverslip which was placed in a 35mm polystyrene tissue-culture dish. Transfections were performed by using the FUGENE 6 transfection reagent (Promega) according to manufacturer’s instructions. Cells were transfected overnight with 1.5 μg per plasmid when the density reached ∼80% confluency. For imaging, coverslips were transferred to FluoroBrite DMEM (Gibco). The spinning disk confocal was performed on an inverted research microscope Eclipse Ti2-E with the Perfect Focus System (Nikon), equipped with a Plan Apo 60× NA 1.40 oil objective, a Crest X-Light V3 spinning disk confocal head (Crest-Optics), a Celesta light engine (Lumencor) as the light source, a Prime 95B 25MM sCMOS camera (Teledyne Photometrics) and controlled by NIS elements AR software (Nikon). A live-cell imaging chamber (H301-Nikon-TI-S-ER, Oko Labs) was equipped to the microscope to provide optimal culture conditions (95% air/5% CO2 atmosphere at 37°C) for cells during imaging. The cell-containing glass coverslip was mounted on a coverslip holder (SC15012, Aireka Cells), which was finally mounted on the microscope.

The fluorescent images for BEAS-2B cells were collected using a 60× oil immersion objective at 2.5 fps (frame per second) for 2 min.

### Single particle tracking and analysis

Single particle tracking (SPT) was performed using a homemade tracking program written in Matlab as previously described^52^, which is based on the standard tracking algorithm^53^. With this advanced SPT algorithm, we were able to obtain the position ***r****(t)* at time *t* for EB1-comets, and their trajectories were constructed from the consecutive images. To characterize the intracellular dynamics of EB1-comets across the whole cell, we first selected the EB1 trajectories that exhibit clear directed motility. This is achieved by computing the mean squared displacements (MSDs), 〈Δ𝒓^!^(𝜏)〉 = 〈(𝒓(𝑡 + 𝜏) − 𝒓(𝑡))^!^〉, and fitted to 〈Δ𝒓^!^(𝜏)〉 = 4𝐷_"##_𝜏^$^ to obtain the effective diffusion coefficient, *Deff*, and the scaling exponent, 𝛼. The scaling exponent was used to classify EB1-comets motion as directed (𝛼 ≥ 1.5)^54^. Polymerization events per minute is defined as the number of directed events detected in 1 minute in individual cells.

### Western blotting

Male flies (24 hours post-injury) were flash frozen in liquid N2 and stored at -80°C. Heads were isolated from fly bodies on dry ice using frozen metal sieves (USA Standard Test Sieve No.25 710µm, Newark Wire Cloth Company). For total protein level analysis, fly heads were homogenized by using Kimble Pellet Pestle Motor and Pellet Pestle in fly lysis buffer containing 50 mM Tris-HCl, pH 8, 150 mM KCl, 2 mM MgCl2, 1 mM EGTA, 0.1% Triton X-100, 1 mM dithiothreitol (DTT), protease inhibitor (1:100, Promega, Madison, WI) and phosphatase inhibitor (1:100, Signma Aldric P5726). Cellular debris was removed by brief centrifugation in a microcentrifuge (BioExpress C-1301-PC) for 20-30 seconds. The samples were heated in the Laemmli SDS sample buffer for 3 minutes at 95°C before electrophoresis. For electrophoresis separation, lysates were resolved using Novex™ Tris-Glycine Mini Protein Gels, 4–12% (Invitrogen, XP04120BOX) in standard Tris/glycine PAGE buffers and transferred to a PVDF membrane (Invitrogen IB24002) using the iBlot 2 Gel Transfer Device (Invitrogen™ IB21001, 20V 4 minutes). Blots were blocked in 5% milk in TBS for 1 hour at 4°C, probed with primary antibodies overnight at 4°C, and fluorescent secondary antibodies for 1 hour at 4C in 5% milk in TBS supplemented with 0.1% tween-20. Primary antibodies were mouse T46 against human tau (1:1000, Invitrogen) and rabbit phosphothreonine (1:500, Cell Signaling). Blots were developed by infrared laser scanner (Odyssey CLx, Licor).

### Statistics

Data are expressed as mean ± s.d. unless specified otherwise. Graphs were created using GraphPad Prism. Statistical tests were performed with two-tailed unpaired Student’s t-test. The statistical details of each experiment can be found in the figure legends.

## Acknowledgements

The authors wish to thank all the members of the Ori-McKenney and McKenney laboratories for their kind help and feedback. This work is supported by the Pew Foundation Scholar Award to K.M.O.-M. and NIH grant 1R35GM124889-06 to R.J.M.

## Author Contributions

CJS. and KMOM conceived the project and designed the experiments. CJS, RM, and KW performed the fly experiments and analysis for Figure 1. CJS, RM, CY, AS, and AF performed the fly experiments and analysis for Figure 2. RM purified the proteins and RM and MFH performed in vitro reconstitution assays and analysis. RM and YS performed the cell experiments and analysis. RM performed all other experiments and analyzed the data. KMOM wrote the manuscript and all authors edited the manuscript.

## Lead contact

Further information and requests for resources and reagents should be directed to and will be fulfilled by the lead contact, Kassandra M. Ori-McKenney.

## Materials availability

Cell lines, fly lines, and plasmids are available upon request from the authors.

## Data and code availability

Data generated and Matlab code used in this study are available upon request.

---

**Supplemental Figure 1.**
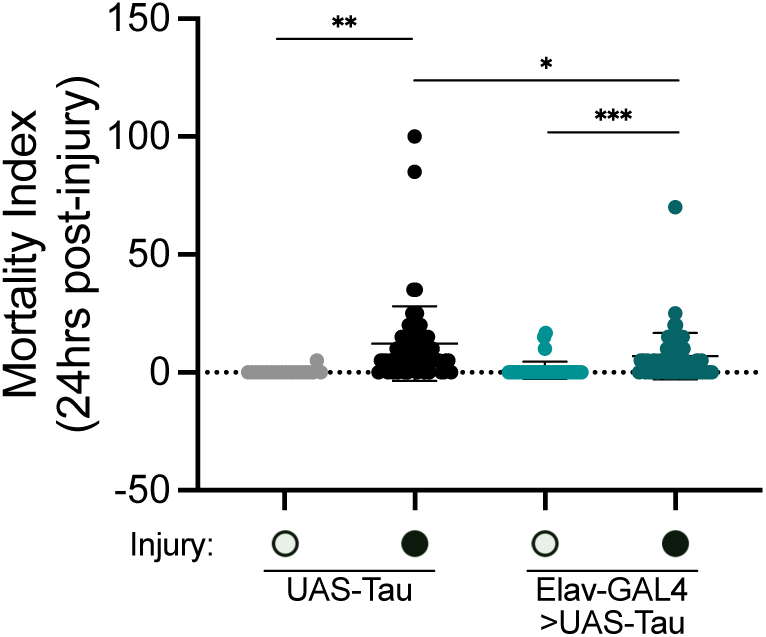
A**d**ministration **of blunt force results in slightly increased mortality 24 hours post-injury.** Quantification of the mortality index of male flies 24 hours post-injury for UAS-Tau sham, UAS-Tau injury, Elav-GAL4>UAS-Tau sham, and Elav-GAL4>UAS-Tau injury conditions (n= 22, 72, 47, and 71 flies, respectively). Two-sided unpaired *t*-tests were used to determine *p*-values. For UAS-Tau sham vs. UAS-Tau injury *p=*0.0011; UAS-Tau sham vs. Elav-GAL4>UAS-Tau sham *p=*0.4042; Elav-GAL4 sham vs Elav-GAL4 injury *p=*0.0002; UAS-Tau injury vs Elav-GAL4 injury *p=*0.0173.

**Supplemental Figure 2.**
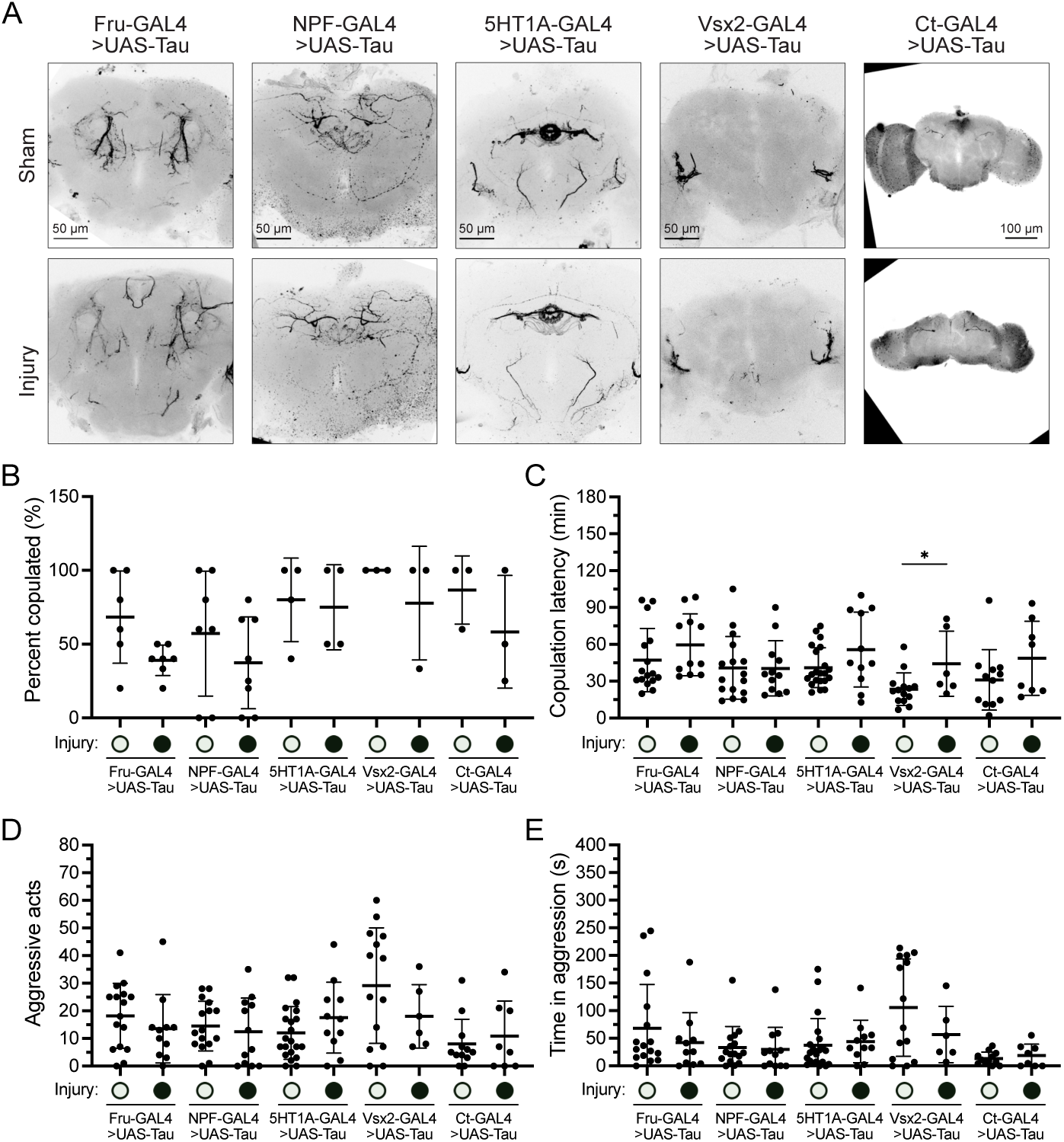
G**A**L4 **lines driving human tau that did not produce significant differences in male fly aggression after injury. A.** Representative max projection images of whole brain immunohistochemistry using an antibody against human tau in Fru-GAL4, NPFGAL4, 5HT1A-GAL4, Vsx2-GAL4, or Ct-GAL4>UAS-Tau male flies for sham and injury conditions. Scale bars: 50 or 100 μm. **BE** GAL4>UAS-Tau male flies for sham and injury conditions. Scale bars: 50 or 100 µm. **B-E.** Competitive courtship assays for sham and injury male flies expressing UAS-Tau under Fru-GAL4 (n=16 and 11 groups, respectively, from 6 independent trials), NPF-GAL4 (n=16 and 12 groups, respectively, from 7 independent trials), 5HT1a-GAL4 (n=21 and 11 groups, respectively, from 4 independent trials), Vsx2-GAL4 (n=14 and 6 groups, respectively, from 3 independent trials), and Ct-GAL4 (n=12 and 8 groups, respectively, from 3 independent trials). **B.** Quantification of percent of flies that mated within 2 hours. Two-sided unpaired *t*-tests were used to calculate *p*-values between sham and injury conditions. For Fru-GAL4>UAS-Tau *p=*0.0385; NPF-GAL4>UAS-Tau *p=*0.3153; 5HT1a-GAL4>UAS-Tau *p=*0.8128; Vsx2- GAL4>UAS-Tau *p=*0.3739; Ct-GAL4>UAS-Tau *p=*0.3332. **C.** Quantification of latency time to copulation. Two-sided unpaired *t*-tests were used to calculate *p*-values between sham and injury conditions. For Fru-GAL4>UAS-Tau *p=*0.2258; NPF-GAL4>UAS-Tau *p=*0.9606; 5HT1a-GAL4>UAS-Tau *p=*0.0826; Vsx2-GAL4>UAS-Tau *p=*0.0296; Ct-GAL4>UAS-Tau *p=*0.1716. **D.** Quantification of total number of aggressive acts exhibited by male flies in the 10-minute window prior to copulation. Two-sided unpaired *t*-tests were used to calculate *p*-values between sham and injury conditions. For Fru-GAL4>UAS-Tau *p=*0.3331; NPF-GAL4>UAS-Tau *p=*0.6064; 5HT1a-GAL4>UAS-Tau *p=*0.1766; Vsx2-GAL4>UAS-Tau *p=*0.2394; Ct-GAL4>UAS-Tau *p=*0.5590. **E.** Quantification of the time spent engaged in aggressive acts by male flies in the 10-minute window prior to copulation. Two-sided unpaired *t*-tests were used to calculate *p*-values between sham and injury conditions. For Fru-GAL4>UAS-Tau *p=*0.3627; NPF-GAL4>UAS-Tau *p=*0.7993; 5HT1a-GAL4>UAS-Tau *p=*0.6944; Vsx2-GAL4>UAS-Tau *p=*0.2240; Ct-GAL4>UAS-Tau *p=*0.4772. All graphs display all datapoints with lines indicating means ± s.d.

**Supplemental Figure 3.**
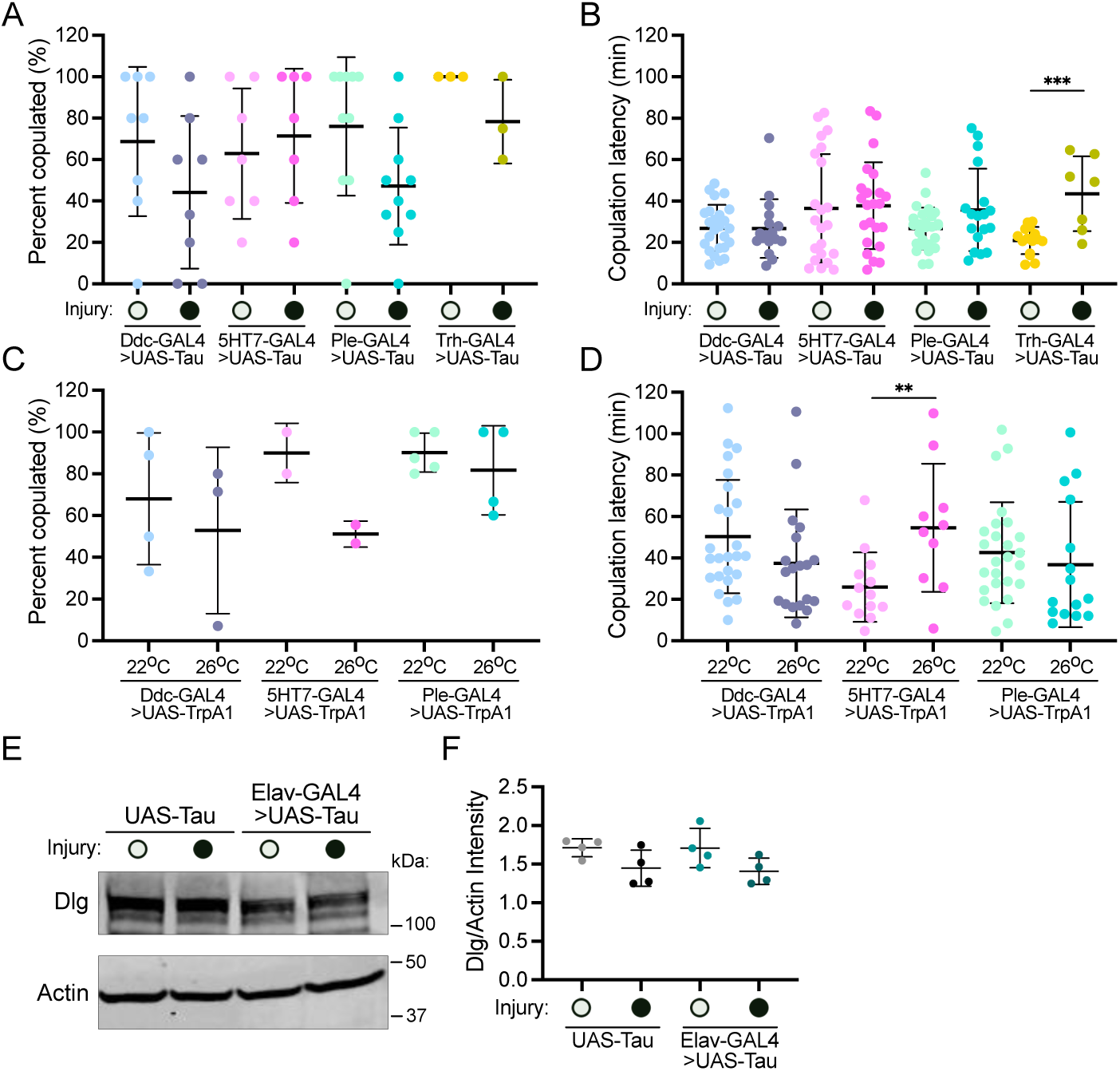
A**s**sociated **analyses of behavioral phenotypes of specific neuronal GAL4 lines. A-B.** Competitive courtship assays for sham and injury male flies expressing UAS-Tau under Ddc-GAL4 (n=25 and 17 groups, respectively, from 8 independent trials), 5HT7-GAL4 (n=22 and 23 groups, respectively, from 7 independent trials), Ple-GAL4 (n=26 and 18 groups, respectively, from 9 independent trials), or Trh-GAL4 (n=15 and 7 groups, respectively, from 3 independent trials). **A.** Quantification of the percent of fly pairs that mated within 2 hours. Two-sided unpaired *t*-tests were used to determine *p*-values between sham and injury conditions. For Ddc-GAL4>UAS-Tau *p=*0.1987; 5HT7-GAL4>UAS-Tau *p=*0.6245; Ple-GAL4>UAS-Tau *p=*0.0518; Trhn-GAL4 *p=*0.1369. **B.** Quantification of latency time to copulation. Two-sided unpaired *t*-tests were used to calculate *p*-values between sham and injury conditions. Ddc-GAL4>UAS-Tau *p=*0.9636; 5HT7-GAL4>UAS-Tau *p=*0.8620; Ple-GAL4>UAS-Tau *p=*0.0563; Trh-GAL4 *p=*0.0003. **C-D.** Competitive courtship assays for male flies expressing the temperature sensitive cation channel UAS-TrpA1 after 1 day of no thermal activation (22 °C) or thermal activation (26 °C) under Ddc-GAL4 (n=25 and 19 groups, respectively, from 4 independent trials), 5HT7-GAL4 (n=13 and 12 groups, respectively, from 2 independent trials), or Ple-GAL4 (n=26 and 15 groups, respectively, from 5 independent trials). **C.** Quantification of the percent of fly pairs that mated within 2 hours. Two-sided unpaired *t*-tests were used to determine *p*-values between temperature conditions. For Ddc-GAL4>UAS-TrpA1 *p=*0.6163; 5HT7-GAL4>UAS-TrpA1 *p=*0.1182; Ple-GAL4>UAS-TrpA1 *p=*0.5004. **D.** Quantification of latency time to copulation upon activation of specific neuronal circuits (26 °C) compared to no activation (22 °C). Two sided unpaired *t-*tests were used to calculate *p*-values between temperature conditions. For Ddc-GAL4>UAS-TrpA1 *p=*0.1202; 5HT7-GAL4>UAS-TrpA1 *p=*0.0095; Ple-GAL4>UAS-TrpA1 *p=*0.5076. **E.** Western blot analysis of lysates generated from the heads of UAS>Tau or Elav-GAL4>UAS-Tau male flies subjected to sham or injury treatment. Blots were probed with antibodies against endogenous Dlg, a post-synaptic marker, and actin. **F.** Quantification of Dlg signal normalized to actin signal. Two-side unpaired *t*-tests were used to calculate *p*-values (n=3 blots). For UAS-Tau sham vs. injury *p* =0.0906; UAS-Tau injury vs. Elav-GAL4>UAS-Tau injury *p* =0.7932; Elav-GAL4>UAS-Tau sham vs. injury *p=*0.0989. All graphs display all datapoints with lines indicating means ± s.d.

**Supplemental Figure 4.**
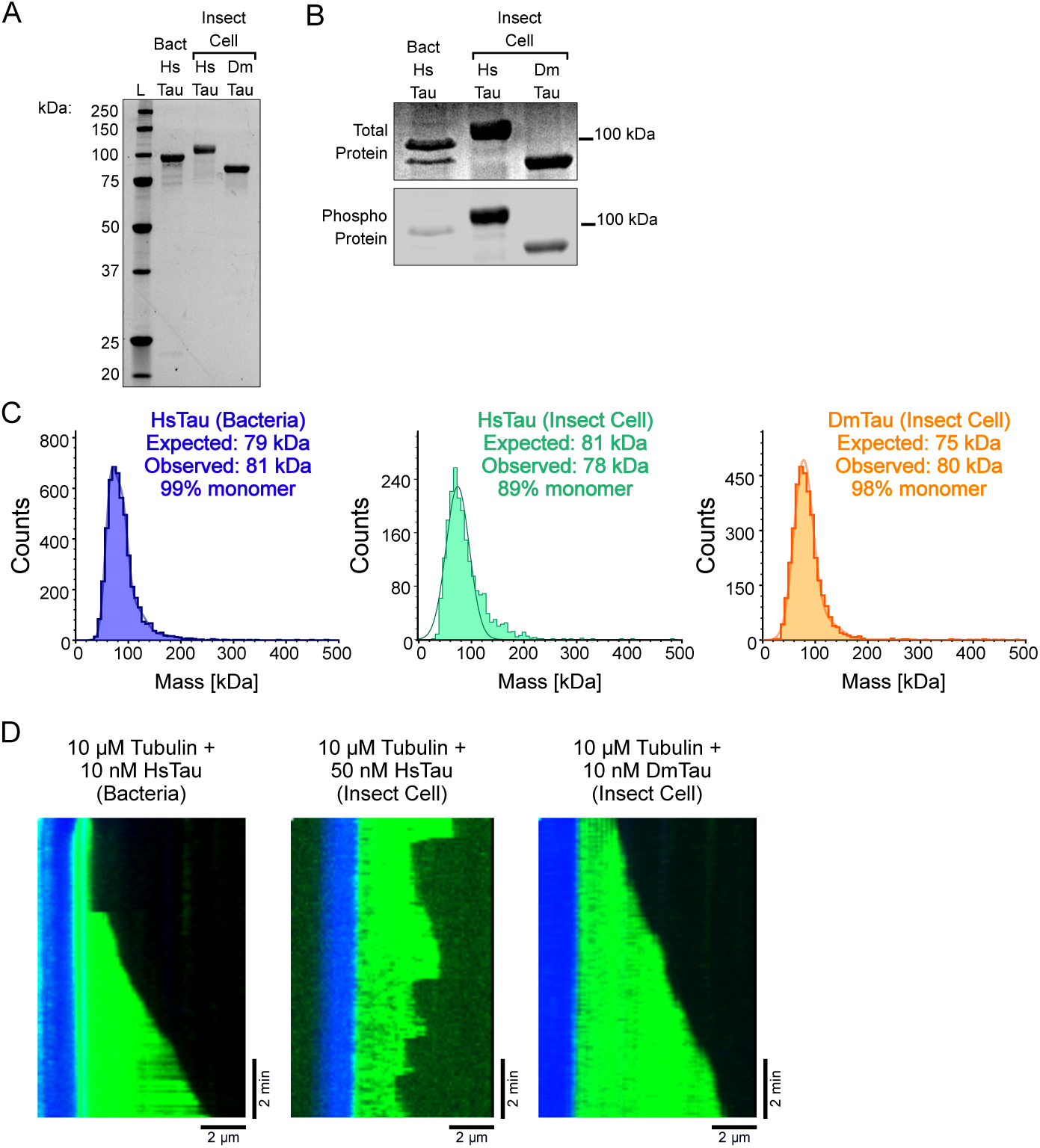
E**x**pression **system dependent phosphorylation and functional properties of tau proteins. A.** SDS-PAGE gels of bacterially expressed sfGFP-HsTau, insect cell expressed sfGFPHsTau, and insect-cell expressed sfGFPDmTau. **B.** SDSPAGE gels of bacterially expressed sfGFP-HsTau, insect cell expressed sfGFPHsTau, and insect-cell expressed sfGFPDmTau stained with SYPRO for total protein (top) or Pro-Q Diamond for phosphorylated protein(bottom) shows significantly more phosphorylation for proteins purified from insect cells compared to bacteria. **C.** Mass photometry of bacterially expressed sfGFP-HsTau, insect cell expressed sfGFP-HsTau, and insect-cell expressed sfGFP-DmTau with indicated observed vs. expected masses. Fits of mean mass values for monomer species, and relative fraction of particles with indicated mass, are shown. **D.** Additional representative kymographs of microtubule dynamics showing the polymerization of 10 μM tubulin (magenta) + 1 mM GTP from GMPCPP seeds (blue) in the absence or presence of human sfGFP-2N4R-tau (HsTau, green) expressed in bacteria or insect cells or *Drosophila melanogaster* sfGFP-tau (DmTau, green) expressed in insect cells at the indicated concentrations. Scale bars: y, 2 min; x, 2 µm.

